# Large Impact of Genetic Data Processing Steps on Stability and Reproducibility of Set-Based Analyses in Genome-Wide Association Studies

**DOI:** 10.1101/2025.07.21.665850

**Authors:** Naishu Kui, Yao Yu, Jaihee Choi, Zachary R. McCaw, Xihao Li, Chad Huff, Ryan Sun

## Abstract

Genome-wide association studies (GWAS) are crucial to human genetics research, yet their stability and reproducibility are often questioned. This work describes, analyzes, and provides tools for overcoming reproducibility challenges in two highly popular components of GWAS: set-based (a) hypothesis testing and (b) effect size estimation. Specifically, we focus on how the set-based natures of (a) and (b) often fuel non-reproducible results due to differences in data processing pipelines that are rarely discussed. First, we describe the processing challenges in a statistical model misspecification framework. Second, we analytically calculate the differences in power and amounts of bias that can arise in (a) and (b), respectively, due to small data processing choices. Third, we provide tools for quantifying and avoiding the data processing obstacles in GWAS. We validate our analytical calculations through a simulation study, and we demonstrate the aforementioned challenges empirically through analysis of a whole-exome sequencing study of pancreatic cancer.

**Author Summary:** The lack of reproducibility and stability in genome-wide association studies (GWAS) have been widely reported. Here, we demonstrate how such reproducibility challenges arise in a common component of GWAS, set-based hypothesis testing and estimation studies. Specifically, we show how minor, seemingly harmless decisions in how scientists prepare their data can lead to major differences in the final conclusions. These data processing steps are rarely reported in detail, further obscuring their importance. Our work precisely measures the impact of data processing steps on power and bias of common modeling approaches. As a partial solution for future GWAS, we also provide an R software package to interact with our results, which can be used to assess the impact of choices at the design stage of GWAS studies. We further analyze a pancreatic cancer dataset using two modern pipelines and show how the pipelines produce very different results for *ATM*, a gene that has been previously linked with pancreatic cancer. Using the tools provided by this work can help significantly improve the reproducibility and stability of set-based results, enhancing the translational potential of GWAS investigations.

## Introduction

Genome-wide association studies (GWAS) are a pillar of modern genetics research, and they continue to be published at high rates [1–5]. In these studies, it is common to perform multiple set-based (a) hypothesis testing and (b) effect size estimation analyses. For example, (a) occurs when associating the outcome against a set of single nucleotide polymorphisms (SNPs) in a gene and (b) occurs when fitting multivariate regression models to estimate polygenic scores or fine-map independent causal effects [6, 7]. However, such analyses can produce highly unstable and non-replicable results due to complex upstream data processing pipelines that are rarely discussed [8–10].

Set-based analysis are ubiquitous in modern GWAS (Supplementary Figure 1) partly because individual SNP association strategies possesses well-known limitations [11, 12].

Compared with individual variant association testing, performing (a) through a set-based method such as the sequence kernel association test (SKAT) [12] offers more power to identify risk factors and can return results that are more interpretable than individual SNP analysis [13, 14]. Such testing is especially useful for rare variant settings but is also widely used for common variants. Additionally, applying (b) to estimate polygenic risk scores or fine-map independent risk SNPs can offer more sophisticated approaches to modeling genetic effects on disease [15].

However, set-based methods applied to massive modern datasets often produce unstable, spurious findings across important use cases, such as cancer studies [16–18]. More specifically, set-based investigations often highlight dozens of putatively important risk factors, yet many of these associations cannot be replicated in follow-up work or even re-analysis of the same cohort [8–10]. The high percentage of unreproducible findings in GWAS is often cited as one of the largest challenges in genetics research.

The major goals of this article are to (i) describe, (ii) analyze, and (iii) provide tools for avoiding data quality obstacles that limit the stability and reproducibility of set-based genetic analyses in modern GWAS. We first (i) describe how seemingly small data cleaning choices -including the choice of annotation database for grouping SNPs into sets, the processing of multi-allelic SNPs, and the selection of set inclusion-exclusion criteria - can lead to large differences in findings. For example, different choices in how to perform regression with multi-allelic SNPs, which make up over 10% of all variants in our data example, can cause large changes in p-values. These issues are characterized quantitatively through a model misspecification framework.

We next (ii) provide analytical calculations for large differences in hypothesis testing power and the large variations in regression coefficient estimates that arise due to the data processing choices and model misspecification. These calculations focus on inference through SKAT - one of the most popular set-based tests - as well as multivariate linear and logistic regression [19, 20]. The analytical calculations are confirmed with simulation to illustrate how results can vary substantially between two studies of the same outcome in the same population.

Finally, we (iii) provide some guidance towards more stable and more replicable analysis. In particular, we introduce an open-source software package, SetDesign, that can be used to design studies and perform sensitivity analysis. We further discuss how the tenets of well-established data science philosophies [21] can be extended to the set-based GWAS setting.

To demonstrate the stability and reproducibility challenges we discuss, we analyze a whole-exome sequencing (WES) study of pancreatic cancer. Pancreatic cancer serves as an useful example phenotype because it is one of the many diseases where sample sizes are still small to moderate [22]. Known genetic risk factors explain only a modest proportion of the estimated heritable risk, and the necessity of set-based analysis is magnified. Pancreatic cancer is also an incredibly deadly disease with urgent need for new risk screening and therapeutic options [23, 24].

Slightly less than half of the pancreatic cancer dataset was previously studied and published [25]. The second stage of the data collection is now complete, and the full study includes 3,360 cases and 4,075 controls. We analyze the full dataset using both the variant-set test for association using annotation information (STAAR) pipeline [26, 27] - a leading modern GWAS package - and the data pipeline of the original study, finding enormous differences even though the two pipelines are applied to the same dataset.

## Materials and Methods

### Overview of Common Data Processing Choices

We briefly describe some of the major choices that modern genetic data pipelines make so that we can model them quantitatively below. These choices often lurk unreported in the background of GWAS analyses, but as we will show, they individually and collectively possess a large impact. The list of factors is not meant to be exhaustive but rather is motivated by empirical observations (Supplementary Figure 1) and is meant to show that our statistical models are sufficiently generalizable.

- *Functional Annotation Database Selection*. Many set-based analyses now form variant sets based on the presumed function of a SNP. For example, the default STAAR pipeline offers the option to filter by promoters, enhancers, and many other groups, which are defined by annotation databases such as FAVOR [28]. However, different annotation databases will offer different groupings [29–31] or disagree on the classifications of a variant. Thus, the choice of database can determine which variants are included or excluded in a set.
- *Multi-allelic SNPs*. Previous studies have estimated that 6-10% of SNPs will be multi-allelic in a sample size of 100,000 subjects [32, 33]. In our pancreatic cancer example, multi-allelic SNPs account for over 10% of all variants. It is possible to both combine all non-reference alleles into a single covariate or consider each non-reference allele separately. That is, if there are three possible alleles at a single location, a regression model could include two separate covariate terms to model this single location.
- *Minor Allele Frequency Filtering*. Some studies perform set-based analyses only using SNPs with minor allele frequencies (MAFs) below a maximum MAF threshold, above a minimum MAF threshold, above a certain minor allele count, or some combination of these criteria. However, minor allele frequencies vary between cohorts, and this issue is especially challenging for rare variants, which can make up around 97% of all variants in large studies [34].
- *Quality Control and Other Filters*. Various other filters, such as genotype quality scores, may be used to include or exclude SNPs from a set. Thresholds for these filters can vary, and therefore such filters may also significantly alter the composition of genotype sets in analysis [35].

A review of recent literature (Supplementary Figure 1) demonstrates that only approximately half of studies (52%) provide any of the above information when describing their set-based analyses. The most common information to provide is minor allele frequency cutoffs for variants, while no papers provide information about how to model multi-allelic SNPs.

### Hypothesis Testing Model

We first consider the effect of different data pipeline choices on (a), hypothesis testing. SKAT and Burden [36] tests are two of the most popular set-based tests. Since Burden is a special case of SKAT-optimal [37] and is much easier to analyze, we focus on SKAT and briefly review its framework here for convenience. Suppose we collect data on *i* = 1, …, *n* subjects, and for each subject we observe a phenotype *y*_*i*_, a *p* × 1 genotype vector **G**_*i*_ = (*G*_*i*,1_, …, *G*_*i,p*_)^*T*^, and a *m* × 1 non-SNP covariate vector

**X**_*i*_ = (*X*_*i*,1_, …, *X*_*i,m*_)^*T*^. For the full sample, we have the *n* × 1 phenotype matrix **y**, a *n* × *p* genotype matrix **G**, and as a *n* × *m* covariate matrix **X**, respectively.

When the outcome is continuous, we assume the data follows the linear model

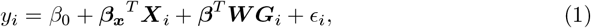

and when the outcome is binary, we assume the data follows the logistic model

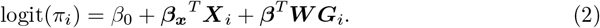

In the SKAT framework, ***β*** is a *p* × 1 random vector of regression coefficients corresponding to the genetic variants, and ***β***_***x***_ is an *m* × 1 vector of fixed regression coefficients corresponding to the non-genetic covariates. The *ϵ*_*i*_ is a random error term with mean 0, and *π*_*i*_ = *E*(*Y*_*i*_ | ***X***_*i*_, ***G***_*i*_) = *P* (*Y*_*i*_ = 1 | ***X***_*i*_, ***G***_*i*_). The pre-specified weight matrix ***W*** is used to upweight variants thought to be more functional. Finally, each element of the random ***β*** is assumed to have mean 0 and variance *τ*. Testing the null hypothesis that there is no association between the phenotype and genotypes is equivalent to testing *H*_0_ : *τ* = 0. Consider first the linear model in Eq (1). The score test statistic is 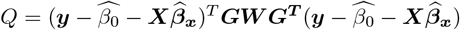 where 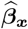 are the non-genetic regression coefficients estimated under the null hypothesis and ***y*** = (*y*_1_, …, *y*_*n*_)^*T*^. When the model is correctly specified, previous work has described how to calculate the power of the test [37].

### Misspecified Model Power

The effect of data processing differences, such as those described in bullet points above, is to create multiple different fitted working regression models. For example, if a set of *p* − 1 variants contains one tri-allelic variant and that variant is represented as one single genetic covariate, then **G**_*i*_ has length *p* − 1 in the fitted models (1) and (2). However, if the tri-allelic variant is represented as two genetic covariates, then **G**_*i*_ would have length *p*. If the two alternative alleles for the tri-allelic variant have different effects, then the smaller model is misspecified.

More precisely, suppose that instead of the true **G**_*i*_, the genotypes are coded as 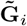, where 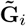 is a *q* × 1 vector of genotypes coded incorrectly. The incorrect vector could be missing some variants due to MAF filtering, it could have a multi-allelic SNP coded as a single variant, or it could be otherwise different from **G**_*i*_. We will assume that *q* < *p*, because the opposite case is generally easier to handle. For instance, if the tri-allelic variant can truly be represented as a single genetic covariate, e.g. both alternate alleles have the same effect, then misspecificying the variant with two covariates simply corresponds to estimating an additional parameter, only lowering power by a small amount.

We write a misspecified working linear model as

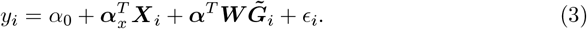

The test statistic is then 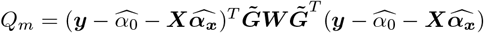 when using the misspecified working model. The 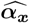 are estimated under the null model with ***α*** = **0**_*q*×1_. Here, for the sake of space, we only outline the power calculation under this misspecified model, and we present the full details in the Supplementary Materials. In brief, the distribution of *Q*_*m*_ is a sum of scaled non-central chi-square random variables 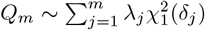, where *m* is the rank of the misspecified genotype correlation matrix. Previous work has shown that such a sum can be approximated well by a single scaled, non-central chi-square random variable through moment matching. The moment matching requires calculation of terms such as 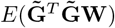 and 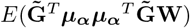, where 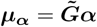. These expressions are complex to evaluate, and we provide full solutions in Supplementary Appendix B for the sake of space.

For binary outcomes, the test statistic of the misspecified model is 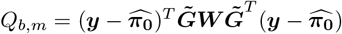 Here 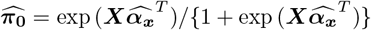 and 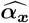 is again estimated under the null model. The test statisti c *Q*_*b,m*_ is still distributed as a sum of noncentral chi-square random variables, 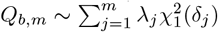. As before, we can perform an analytic power calculation, but the quantities are complex and difficult to display. We provide the full forms in the Supplementary Appendix C.

Our open-source R package `SetDesign` provides analytic solutions to calculate power under misspecified models that may arise from different data processing pipelines. These equations are provided for both linear and logistic regression models. To the best of our knowledge, this is the first time that all terms are provided analytically, instead of relying on simulations or other computationally expensive approaches [38]. Because the implementation in `SetDesign` does not require sample genotypes or bootstrapping the full model, power can be easily calculated for many different scenarios at once. Thus, our software can be used at study design stages to study the impact of potential data processing choices.

The two main novelties of this work compared to previous literature are that (1) it evaluates the potentially-misspecified working model and (2) it is much simpler and more computationally efficient to perform since no actual genotype data is needed. The second feature is a significant advantage over existing tools [39], which generally require input of full genotype matrices to bootstrap the regression model. Such an approach is impractical for modern datasets with massive genotype matrices and many rare variants that would be difficult to extract or simulate.

### Bias in Estimated Effect Sizes

We next consider the bias of (b) effect size estimation under multivariate regression models that are another popular type of set-based study in GWAS. In both polygenic risk score (PRS) and fine-mapping studies, researchers will again fit regression models similar to equations (1) and (2). However instead of testing for association, focus lies in estimates of each SNP’s regression coefficients, which will either be used as score weights in PRS or to justify the existence of multiple independent signals in fine-mapping.

Additionally, in these regression models, all coefficients are treated as fixed parameters. For continuous outcomes, we assume again that model (1) is the true model, except that we suppose ***β*** is now fixed instead of random, so that we can study the bias when it is estimated. In this conventional fixed effects linear regression model, the *m* + *q* + 1 score equations for estimating *α*_0_, ***α***_*x*_, and ***α*** under misspecified working model (3) are proportional to

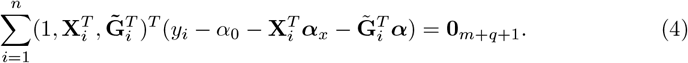

The asymptotic limit ***α*** of the maximum likelihood estimate ***α*** can be found by taking the expectation of the left hand side of equation (4) and solving for ***α***. The full solutions are available in closed form. We provide these in the Supplementary Materials (see Supplementary Appendix D) for sake of space. Our R package SetDesign also provides the solutions to these equations for public use.

As a simple example of how misspecification can significantly alter estimated effect sizes, consider the true model of equation (1) where there are no additional covariates 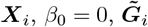 is just one tri-allelic SNP coded as a single genotype, and ***G***_*i*_ is the tri-allelic SNP coded as two genotypes, with both alternative alleles being rare and thus almost independent. Let the effect sizes of the alternative alleles be *β*_1_ and *β*_2_, and let the misspecified effect be *α*. Denote by Var(*G*_1_) and Var(*G*_2_) the variance of binomial random variables with size 2 and probability parameters equal to the minor allele frequencies of the two alternative alleles. Then we have

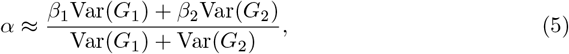

and *α* can be close to zero if *β*_1_ and *β*_2_ are of similar magnitudes but opposite directions. Thus the variant is wrongly estimated to possess almost no effect even though it may be quite important.

For the logistic regression case, the score equations when all regression coefficients are fixed are proportional to

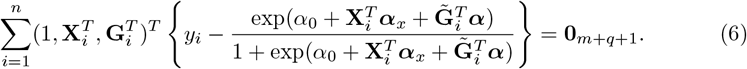

In general there are not closed form solutions similar to the above. However, we can solve them numerically. We provide these numerical solutions in our R package.

### Simulation Studies

We conducted an extensive set of simulations to demonstrate the magnitude of challenges that misspecification poses. For each set of simulations, we consider a set of 50 SNPs. Each SNP is given a minor allele frequency of 5% if they are correlated with other SNPs or 1% in the uncorrelated setting, and we consider a sample size of *n* = 2, 000 individuals.

When the outcome is continuous, the true model is *y*_*i*_ = *β*_0_ + *G*_*i*,49_*β*_49_ + *G*_*i*,50_*β*_50_ + *ϵ*, where *ϵ* ∼ *N* (0, 1). We then fit the misspecified working model 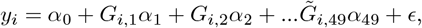, where 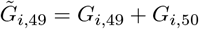. This situation mimics the setting where the last variant in the set is a tri-allelic SNP but we treat it is a bi-allelic SNP and simply sum the number of non-reference alleles. It is also similar to settings where SNPs may be left in or out of sets based on annotations or other filters. When the outcome is binary, the true model is logit(*π*_*i*_) = *β*_0_ + *G*_*i*,49_*β*_49_ + *G*_*i*,50_*β*_50_, and we fit the model 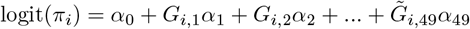, where again 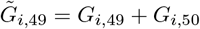.We performed all testing at the 0.05 level.

### Pancreatic Cancer Whole-Exome Sequencing Data

Our work is motivated by a WES study of pancreatic cancer. The dataset includes 3,360 cases and 4,075 controls of European ancestry. The data generation details were previously described [25], and a previous gene-based association study using less than half of the dataset emphasized *ATM* as the main finding. The *ATM* gene includes 3,832 positions with variants in our dataset. In the original analysis, 215 of these positions were analyzed to associate rare coding mutations with pancreatic cancer. Here we list four major differences between the original, reasonable, analysis pipeline and the STAAR default pipeline for rare variant gene-based hypothesis testing. Note that just considering these four factors allows for 2^4^ = 16 different combinations of data cleaning choices.

- The original analysis used all SNPs with MAF less than 0.005 while the STAAR default is a 0.01 threshold.
- The original analysis considered all single and multi-nucleotide variants, while the STAAR default is single nucleotide variants only.
- The original analysis considered each non-reference allele (of a multi-allelic variant) separately, while an alternative is to group all non-reference alleles into a single alternative allele.
- The original analysis formed SNP sets using all coding variants, which were defined as stop-gain, stop-loss, splicing, synonymous, nonsynonymous, frameshift, and non-frameshift insertion and deletion variants. The STAARpipeline software (prior to v0.9.7) does not include the last two categories in its definition of coding variants.

We performed analysis using both the original analysis settings [25] and STAAR default settings, which are the opposite of the original analysis for each choice above. When contrasting p-values between the new and original data pipelines, we reported final STAAR p-values (which is a meta-analysis including the unweighted SKAT p-values and other tests) when ranking genes to mimic a practical analysis; the STAAR p-value is generally very similar to the unweighted SKAT p-value. The same non-genetic covariates were used in all models. These choices were made to facilitate balanced comparisons between the different settings.

## Results

### Simulation Studies

In Fig 1, we first demonstrate the concordance between the analytic power calculations and simulated results. We can see that the analytic calculations match the empirical power very closely. This means that even though the calculations use asymptotic arguments, they are still reliable in practical situations. In Fig 1A-D we consider fixing *β*_49_ at four different values and then varying the value of *β*_50_ in the opposite direction with uncorrelated SNPs. When the magnitude of *β*_49_ is small in Fig 1A-B, the model misspecification does not affect the power of the test much. However, when the magnitude of *β*_49_ is larger in Fig 1C-D, the misspecification can have a large effect. When *β*_50_ is in the exact opposite direction of *β*_49_ and both have large magnitudes, the power of testing under the misspecified model can be close to 0, whereas it would be close to 1 if the model were correctly specified. These findings make intuitive sense since large effects in opposing directions would be expected to cancel each other out when the multiple alternative alleles with similar MAFs are combined.

**Fig 1.**
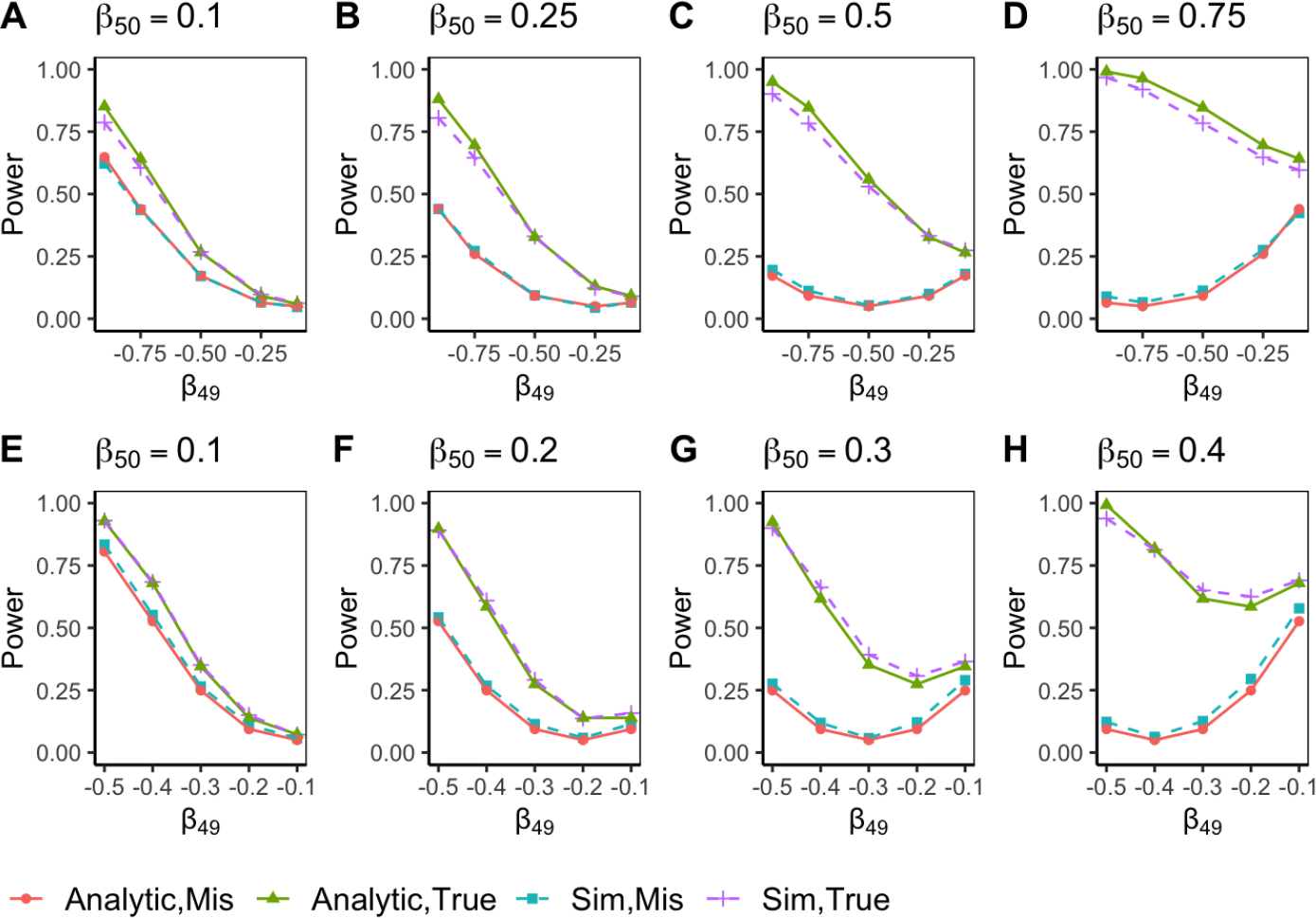
Power simulations for linear outcomes across varying effect sizes, with sample size = 2,000. Simulations were performed using 50 uncorrelated SNPs with MAF = 0.01 (panels A–D) and 50 correlated SNPs with MAF = 0.05 (panels E–H). *Analytic, Mis* represents analytical power for the misspecified model, where two alternative alleles of a triallelic SNP are combined. *Analytic, True* represents analytically derived power for the true model, where alternative alleles at the multi-allelic position are treated as separate variants. *Sim, Mis* represents the simulated power for the misspecified model; *Sim, True* represents the simulated power for the true model. Note that the analytical calculations (solid lines) and simulated powers (dashed lines) align very closely.

In Fig 1E-H, we increase the minor allele frequencies of the SNPs so that it is possible to add some correlation to the genotype set. Here the MAF is 0.05 for all SNPs, and we use an exchangeable correlation structure with common correlation coefficient *ρ* = 0.15. We lower the effect size to show the power difference. We can see that the power loss due the misspecification is in general less in Figure 1E-F compared to Figure 1A-B. The overall power is also generally higher (Fig 1G-H) than the uncorrelated setting. This occurs because the correlation spreads effects [13] to the non-causal SNPs, and thus there is more signal overall to detect. However, when *β*_49_ and *β*_50_ have the same magnitude but opposite directions, there can still be extreme power loss.

In Fig 2A-D we consider the logistic regression setting with uncorrelated SNPs and MAFs of 0.01 again. Here the effect of misspecification is more muted compared to the linear regression setting. One reason is that the effects of misspecification are much more complex in logistic regression models [40]. However, we can see that one commonality with the linear regression setting is that the differences in power rise as the magnitudes of the causal effects rise.

**Fig 2.**
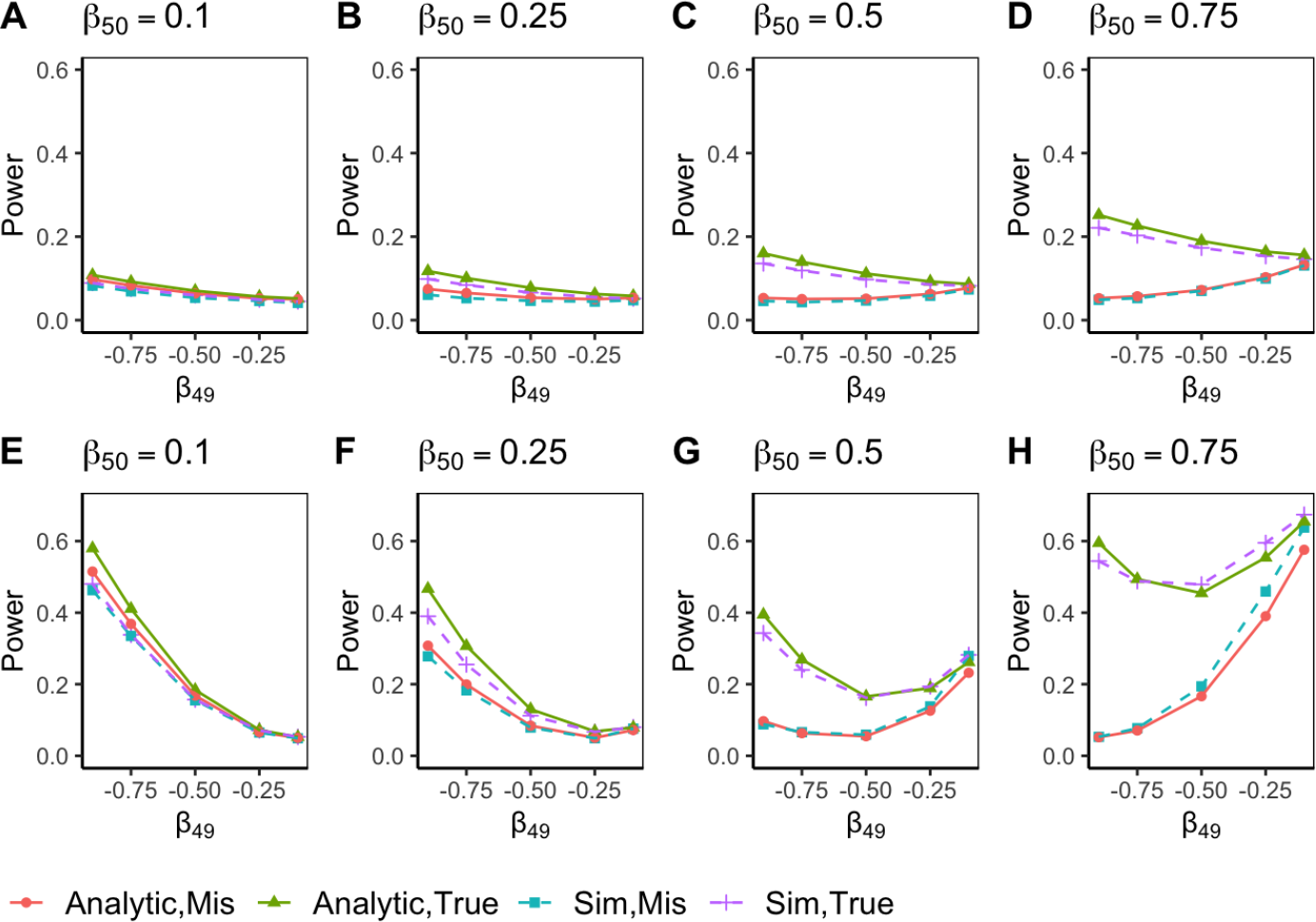
Power simulations for binary outcomes across varying effect sizes, with sample size = 2,000. Simulations were performed using 50 uncorrelated SNPs with MAF = 0.01 (panels A–D) and 50 correlated SNPs with MAF = 0.05 (panels E–H). *Analytic, Mis* represents analytical power for the misspecified model, where alternative alleles are combined. *Analytic, True* represents analytically derived power for the true mode, where alternative alleles at multi-allelic positions are treated as separate variants. *Sim, Mis* represents the simulated power for the misspecified model; *Sim, True* represents the simulated power for the true model.

Finally, Fig 2E-H is the logistic regression analog of Figure 1E-H, where the MAF is increased to 0.05 and there is an exchangeable correlation structure with *ρ* = 0.15. We can see that again there is more power overall. Additionally, at times, the power of the misspecified model can be similar to the power of the correctly specified model. This behavior is a reminder that data processing differences do not always lead to negative results, but rather, other factors such as effect sizes also play a large role. Our software can be used to perform a variety of sensitivity analyses so that the impacts of different data pipeline choices can be studied.

While power loss can be almost total when effects are in opposite directions, there can be significantly negative impacts on power across a variety of other settings as well. Thus, large disparities in p-values across studies of the same outcome in the same population can easily occur due to small differences in data processing that affect only a single variant, as demonstrated above. Further examples can be found in the analysis of the pancreatic cancer data presented below.

For sake of space, simulation studies for bias in estimating regression coefficients are presented in Supplementary Appendix E. In these simulations, we show that our analytic calculations for the bias of the misspecified regression coefficients matches very closely with the empirical simulated bias. We note that the estimated misspecified regression coefficient can be very different from the true effect sizes.

### Pancreatic Cancer Whole-Exome Sequencing Data

Fig 3 and Table 1show differences in gene-based p-values between the original half-cohort analysis pipeline [25] and the default STAAR pipeline gene-based analysis. We can immediately see there are some very large differences. For example, Figure 3 shows how the p-value of *ATM* can vary depending on the data cleaning steps used.

**Table 1.** Genes that make the largest jump or drop in final STAAR significance ranking when switching between original data pipeline and STAAR default pipeline. The final STAAR p-value is a meta-analysis combining the unweighted SKAT p-value with other tests.

| Gene | Data Pipeline Settings | Significance Ranking | #Variants | SKAT P-value | STAAR P-value |
| --- | --- | --- | --- | --- | --- |
| BCO1 | MAF 0.005, SNV, more coding, multi-allelic separated (original) | 16 | 51 | 0.00041 | 0.0014 |
| BCO1 | MAF 0.01, SNV + MNV, fewer coding, multi-allelic combined (STAAR default) | 3330 | 35 | 0.24 | 0.21 |
| ULK1 | MAF 0.005, SNV, more coding, multi-allelic separated (original) | 3745 | 65 | 0.19 | 0.22 |
| ULK1 | MAF 0.01, SNV + MNV, fewer coding, multi-allelic combined (STAAR default) | 1 | 49 | 1.1e-12 | 2.6e-12 |

**Fig 3.**
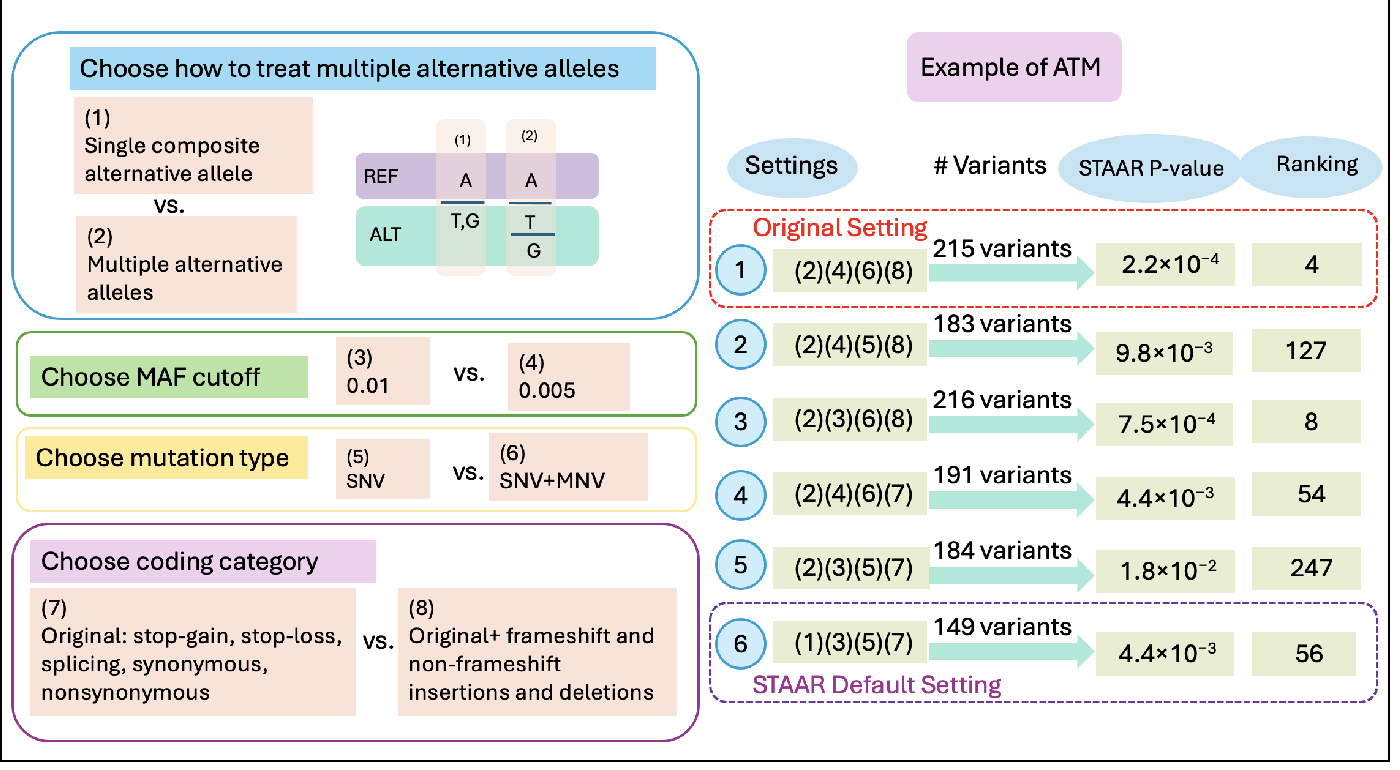
Differences in original publication and default STAAR analysis of pancreatic cancer dataset. On the left we list the four data processing steps that are different, and on the right we show how different combinations of these steps contribute to highly different SKAT and STAAR p-values for the *ATM* gene. The four differences on the left are: treatment of multi-allelic variants, MAF cutoff, types of mutations (single nucleotide vs. multi-nucleotide variant), and types of variants classified as coding. On the right, the original analysis (1) and STAAR default (6) settings rank *ATM* as the fourth most significant and 56th most significant gene, respectively. We also show four additional combinations (2)-(5) of data processing choices that could plausibly be made and the varying p-values they produce. Note that the final STAAR p-value is a meta-analysis combining the unweighted SKAT p-value with other tests.

*ATM* was the most strongly highlighted gene in the original publication on the first half of this dataset, based in part on the 4th most significant p-value, as well as other factors such as annotation data. Germline *ATM* variants have been linked with pancreatic cancer in a variety of other settings as well [41]. However, when using the STAAR default pipeline, the *ATM* p-value ranking drops to 56. In other words, a researcher attempting to reproduce the findings of the original publication would have a difficult time if they made the reasonable choice of using the STAAR default pipeline.

Other genes, such as *BCO1* and *ULK1*, can change from very highly associated with pancreatic cancer to not associated at all, or vice versa, depending on the pipeline choices made. We can see in Table 1 that *BCO1* is the 16th most significant gene when using the original analysis pipeline, but it becomes the 3330th most significant gene under the STAAR default settings. In contrast, *ULK1* is the most significant gene under the STAAR default settings, but it is the 3745th most significant gene in the original analysis pipeline.

Fig 4 shows the genome-wide gene significance profiles through Manhattan plots under different data pipeline choices (more details available in Supplementary Appendix F and G). We can see that some genes do hold relatively similar p-values across different settings, such as *AADACL3*. However, there are also clear differences in the shapes of the plots across most chromosomes. Thus, there is a substantial impact of data processing choices on the results of the gene-centric analysis across all genes, not just the most significant findings.

**Fig 4.**
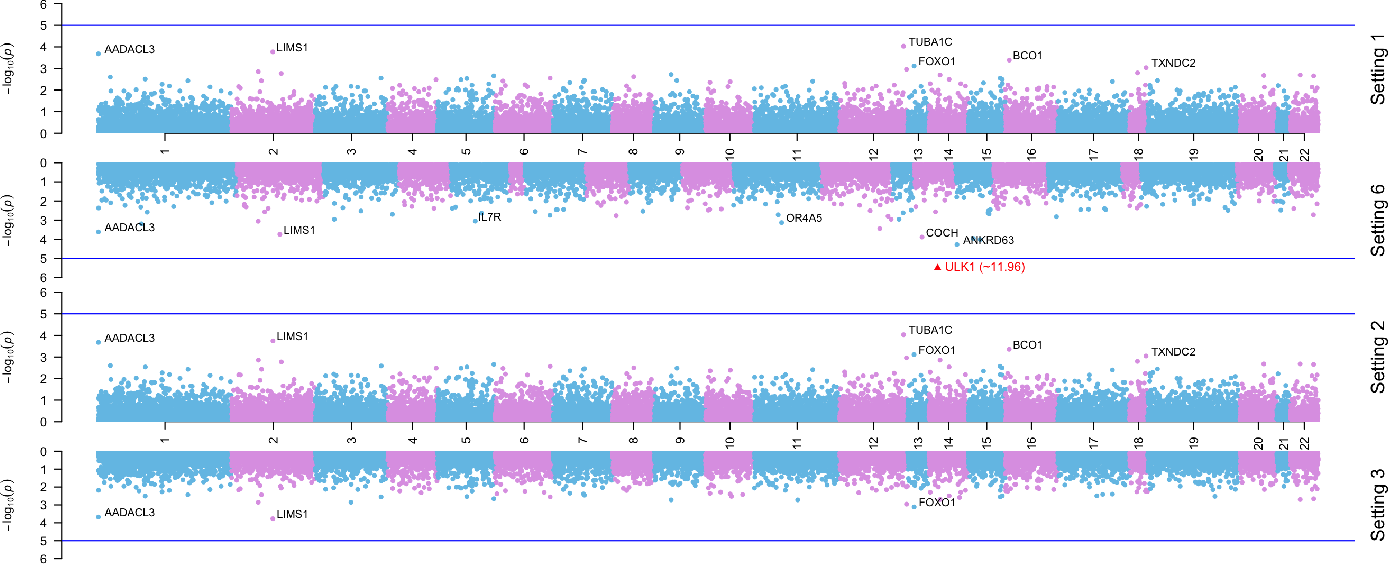
Manhattan plots of gene-based association with pancreatic cancer under settings 1, 2, 3, and 6 (setting numbers from Fig 3). Unweighted SKAT p-values are shown. Setting 1 uses the original publication pipeline while setting 6 is the STAAR default. Blue horizontal line corresponds to *p* = 10^−5^. Some gene p-values (e.g. *AADACL3*) are relatively similar across different settings, while other genes (e.g. *ULK1*) vary wildly.

Finally, Table 2 presents the results of multivariate regression analyses for the top five SNPs in *BCO1* across different settings. *BCO1* is one of the most significant genes in the original analysis (Table 1), and it holds multiple individual SNPs with small p-values. It is reasonable to assume that follow-up PRS construction or fine-mapping would fit regression models with the *BCO1* SNPs. For simplicity and sake of presentation, we consider fitting a standard GWAS logistic regression model that includes the top five SNPs in the gene, which all have *p* < 0.05 in the multivariate model. One of these five SNPs, rs119478057, is a tri-allelic SNP. We fit the model treating rs119478057 as a single term (as in the default STAAR analysis) and then also as two terms, one for each different non-reference allele (as in the original analysis of this dataset). We can see that the estimated regression coefficient changes by about 14%. Given the large number of multi-allelic SNPs in this dataset, such differences in estimated regression coefficients could, for example, significantly affect PRS performance.

**Table 2.** Multivariate regression model including top 5 SNPs in *BCO1* and same non-genetic covariates as original analysis of this dataset. The third row (rs119478057) is a multi-allelic SNP and is modeled with two terms in the original analysis. Only one of those two terms shows significant association. We only show the coefficient and p-value for the significant term.

| rsID | Position | Original Setting |  | STAAR Default Setting |  |
| --- | --- | --- | --- | --- | --- |
|  |  | Coefficient | P-value | Coefficient | P-value |
| rs200504527 | 81285639 | 2.000 | 0.009 | 2.000 | 0.009 |
| rs143003259 | 81267970 | -2.090 | 0.009 | -2.089 | 0.009 |
| rs119478057 | 81264677 | -1.128 | 0.015 | -0.969 | 0.026 |
| rs141202792 | 81270286 | -1.263 | 0.023 | -1.263 | 0.023 |
| rs147813781 | 81264649 | 0.811 | 0.032 | 0.811 | 0.032 |

## Discussion

As large-scale genetic data becomes increasingly accessible and more GWAS are conducted, increasing the stability and reproducibility of results across studies becomes critical. Our research work highlights a key but overlooked challenge in GWAS: the importance of data processing steps in set-based analysis. Studies of the same phenotypes that do not use the same processing pipelines can encounter significant challenges in replicating findings; such a replication crisis is commonly noted as one of the most important problems facing GWAS today.

Our work provides precise analytical calculations for the loss in power and the bias that can arise due to differences in data processing. Specifically, we study different processing choices under a model misspecification framework, considering both linear and logistic regression models. For set-based inference, we demonstrate that data processing choices such as mistakenly combining the effects of a multi-allelic variant can cause almost 100% loss in power. In the estimation setting, we show that multivariate regression models can produce highly biased estimates of effect sizes when pipeline choices lead to misspecification.

When performing gene-based association analysis, we find that common, defensible choices for cleaning genotype data can cause genes such as *ATM* to be either highly associated or not associated at all with pancreatic cancer. Previous studies have both replicated and not replicated the *ATM* association [41], and our work shows that both findings can easily be made even within the same dataset depending on how the gene set is defined. We also demonstrate that regression coefficient estimates at top genes like *BCO1* vary substantially when data are processed differently.

To improve the robustness and reliability of future work, we recommend use of our open-source R package SetDesign, which allows researchers to perform sensitivity analyses on their data pipelines. For example, users can perform quick calculations about power differences that may arise when selecting different MAF thresholds for gene-based testing. These calculations can be performed before any data is collected, thanks to our analytical model misspecification framework.

It should be noted that other, non-statistical approaches could help improve the stability and reproducibility of set-based results. For example, utilizing data science best practices such as the Predictability, Computation, Stability (PCS) principles [21] is also recommended. Briefly, PCS emphasizes publication of both workflows and documentation, to clearly identify important data processing choices. Our R package can be used in conjunction with PCS and other frameworks at the design stage of studies to assess the potential impact of different data pipeline decisions.

